# Integrating simultaneous motor imagery and spatial attention for EEG-BCI control

**DOI:** 10.1101/2023.02.20.529307

**Authors:** Dylan Forenzo, Yixuan Liu, Jeehyun Kim, Yidan Ding, Taehyung Yoon, Bin He

## Abstract

Objective: EEG-based brain-computer interfaces (BCI) are non-invasive approaches for replacing or restoring motor functions in impaired patients, and direct brain-to-device communication in the general population. Motor imagery (MI) is one of the most used BCI paradigms, but its performance varies across individuals and certain users require substantial training to develop control. In this study, we propose to integrate a MI paradigm simultaneously with a recently proposed Overt Spatial Attention (OSA) paradigm, to accomplish BCI control. Methods: We evaluated a cohort of 25 human subjects’ ability to control a virtual cursor in one- and two-dimensions over 5 BCI sessions. The subjects used 5 different BCI paradigms: MI alone, OSA alone, MI and OSA simultaneously towards the same target (MI+OSA), and MI for one axis while OSA controls the other (MI/OSA and OSA/MI). Results: Our results show that MI+OSA reached the highest average online performance in 2D tasks at 49% Percent Valid Correct (PVC), statistically outperforms MI alone (42%), and was higher, but not statistically significant, than OSA alone (45%). MI+OSA had a similar performance to each subject’s best individual method between MI alone and OSA alone (50%) and 9 subjects reached their highest average BCI performance using MI+OSA. Conclusion: Integrating MI and OSA leads to improved performance over MI alone at the group level and is the best BCI paradigm option for some subjects. Significance: This work proposes a new BCI control paradigm that integrates two existing paradigms and demonstrates its value by showing that it can improve users’ BCI performance.

## I. INTRODUCTION

RAIN-computer interfaces (BCI) enable control of an external computer or device by interpreting users’ intents directly from their brain signals. By bypassing the need for muscle activity or speech, BCIs have the potential to replace or restore motor functions for people with motor impairments or even enhance the capabilities of healthy individuals [1]. Already these devices are being used to help some motor-impaired patients interact with the outside world by moving, and even feeling through, tools like robotics arms [2]–[4].

There are two main categories of BCI devices: invasive and non-invasive, depending on the method used to acquire the neural signals. Non-invasive methods allow the device to be used without the user having to go through any invasive medical procedures, although they also have lower signal-to-noise ratios (SNR). Among non-invasive devices, EEG-based BCIs are the most commonly studied due to the low cost, ease of recruiting subjects, and high temporal resolution of the EEG signal [1], [5]. However, while EEG based BCIs have achieved control for complex applications such as control of a wheel chair[6], a virtual [7], [8] and physical [9] quadcopter, a robotic arm [3], [10]), and speech decoding [11], these devices currently have few clinical applications due to low SNR and limited performance.

A focus of the non-invasive BCI field is to improve users’ control of these devices. More reliable performance can make BCIs easier for potential subjects to use and can allow them to perform more complex tasks, which will expand their potential applications. Advancements in the components of BCI systems, such as new recording hardware, signal processing techniques, and training methods, have improved users’ performance in recent years [3], [12]–[22]. However, even with these technological advancements, EEG-BCI performance is limited by the inter-subject variability and low SNR of the scalp-recorded EEG signals produced by the users.

One of the most important components of a BCI system is the control paradigm, or the task that the user performs to control the BCI system. Since the goal of a BCI device is for the user to be able to control an external computer or device using their brain recordings, a successful BCI control paradigm must allow the user to predictably change, or modulate, their neural signals. If the user can modulate their brain activity predictably and consistently, the BCI system will be able to decode their intentions more effectively, leading to better BCI performance. Because of this, the choice of control paradigm has important implications for both the performance and possible applications of a BCI system.

Several different control paradigms have been established for EEG-BCI systems. Some of the well-established methods include the P300 signal, Steady-State Visually Evoked Potential (SSVEP), and Motor imagery [1]. Some of these methods, such as Motor Imagery, are endogenous methods, meaning that the stimulus for modulating the user’s brain activity is created voluntarily by the user. Other methods, such as P300 and SSVEP, are exogenous and require an external stimulus that is not controlled by the user. While non-volitional methods can produce strong and reliable neural signals, they can only be used in the presence of an external stimulus. Among the available control paradigms, this study focuses on two techniques in particular: Motor Imagery [23] and Overt Spatial Attention [24], both of which are endogenous methods.

Motor imagery (MI) is one of the most common methods for controlling non-invasive BCIs. This technique usually involves the user imagining the feeling or sensation of moving a body part to produce neural signals that are similar to those produced by actual movement (see the top two rows of Figure 1) [23]. MI has been used extensively for EEG BCIs, including systems for one, two, and even three dimensional control of virtual objects [7], [8], [25], [26], robotic arms [10], [27], and wheelchairs [6]. While there are several advantages of MI over other established paradigms [28], the strength and location of the signals produced by this method vary across different users. Potential BCI users have a wide range of performance using MI, ranging from near perfect control to BCI illiteracy (unable to gain any meaningful control of the BCI system) [29]. Subjects are known to improve MI performance over time through learning, but this can take many sessions of training and even then, some subjects are still not able to produce effective control [20], [28], [30].

**Figure 1.**
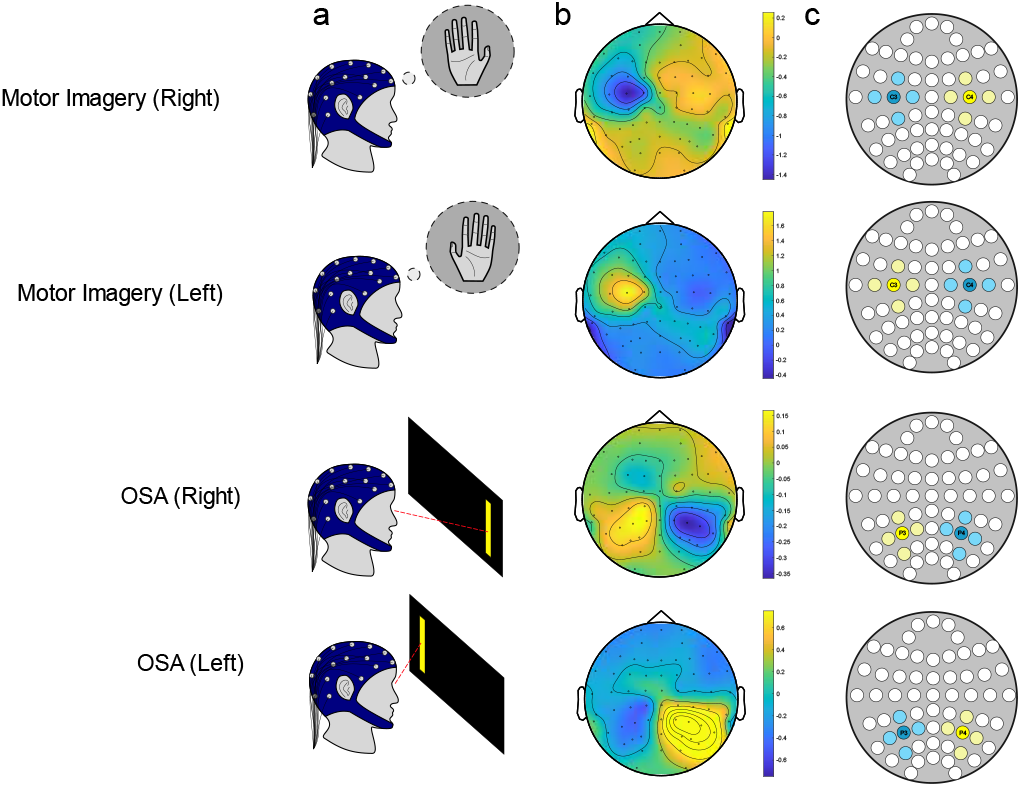
Brain computer interface control techniques. A) The subject is asked to perform a task to control the BCI cursor. Either Motor Imagery (Top) where the subject imagines the feeling or sensation of moving their left or right hand, or OSA (Bottom) where the subject looks and focuses on the target. B) The BCI tasks cause a change in the alpha power (8-13 Hz) of the EEG signal in certain cortical areas depending on the task C) The relative alpha power in certain electrodes used to control the BCI cursor. The specific electrodes used depend on the control method.

Overt spatial attention (OSA) is another, recently proposed, BCI control paradigm. This method involves the user looking at a target and focusing their visual attention on it (Figure 1 bottom) [24]. While this method does require the use of eye muscles to look at a target, it still bypasses the need for other muscle activation and has been shown to produce similar performance as MI in subjects with little to no BCI experience. The original OSA study found no significant changes in OSA performance over 3 sessions of BCI and, to our knowledge, there have been no studies investigating the effects of BCI training or learning on OSA performance over more than 3 sessions. Due to the simplicity of the OSA paradigm and the intuitive nature of looking and focusing on a target, there may be no substantial learning effect for OSA BCI, but more evidence is needed on this subject. Although both MI and OSA have been shown to produce similar average BCI performance across subjects, the advantage to having both paradigms available is that individual subjects may perform much better with one paradigm over the other. In addition, since these two tasks activate different cortical areas, they can potentially be performed at the same time.

This study investigates the effects of integrating two independent BCI control methods simultaneously to control a virtual computer cursor. Previous studies have combined BCI control methods before, including combinations of exogenous methods such as SSVEP and P300 [31]–[33], or even combinations of EEG with other recording modalities [34]. Here, we combine two endogenous methods simultaneously towards the same goal. The main question this work seeks to answer is whether combining these two different control paradigms could produce a stronger overall control signal that would lead to better BCI performance than either individual method, or if their performance would remain somewhere in between the two. The results from this work show that, using a simple online linear classifier, subjects reached the highest average 2D performance with the combined MI and OSA paradigm. This combined method performed substantially better than MI alone, and had a higher mean performance, but not statistically significant, above OSA alone. This method also reached about the same online performance as using the best individual method for each subject at the group level (either MI or OSA alone). In addition, some subjects in the cohort were able to achieve their best overall BCI control using the combined method. Although it is limited to a subset of the study population, this represents a novel improvement in BCI performance attained by combining two established control methods.

In addition to showing the promise of combining BCI tasks, the results here emphasize the differences in individual BCI users. While both tasks led to similar performance at the group level, some subjects performed better substantially better using MI rather than OSA, and others using OSA rather than MI. Similarly, some subjects benefitted from combining MI and OSA, while others performed much worse than they could have if they used one of the individual methods instead. These conclusions highlight the need to design BCI systems around the user’s strengths and show the benefit of developing more control task options, since what has been shown to work best for some users does not work well for others. In the end, improving individual users’ BCI performance will lead to increased consistency and reliability of non-invasive BCI devices, and can expand their potential clinical uses.

## II. MATERIALS AND METHODS

Figure 1 illustrates how both MI and OSA produce signals that can be detected from EEG systems. In both cases, the subject voluntarily performs the task with a specific intent. In the case of virtual cursor control, this can be performing MI or OSA to move the cursor to the left or right (Figure 1a). This task causes a change in certain frequency ranges of the EEG signal, particularly around the 8-13 Hz range known as the alpha power of the signal (Figure 1b). By focusing on these frequency ranges in electrodes over specific brain areas, BCI systems are able to detect these power changes and respond accordingly (Figure 1c). Different BCI tasks cause changes in various frequency bands and EEG electrodes, so most decoders are modified to fit different tasks. Because control methods produce neural signals in different areas of the cortex, combining them may result in more potential electrodes with decodable features that could produce a more reliable output signal, even when one of the individual methods fails.

### A. Subject Recruitment

Twenty-five (average age: 25.5, 24 right-handed, 15 male) healthy human subjects were recruited for 5 sessions of BCI experiments. Subjects were recruited through fliers placed around Carnegie Mellon University campus in Pittsburgh, PA, and every subject that was recruited completed all 5 sessions. Each subject provided written consent to the protocol approved by the Institutional Review Board of Carnegie Mellon University. The subjects had varying levels of BCI experience, from no prior experience to having participated in several previous BCI studies.

### B. Data Collection and Online Processing

EEG data were collected using the Neuroscan 64 channel Quik-Cap with SynAmps 2/RT amplifier system (Compumedics Neuroscan) with electrodes in an extended version of the international 10/20 system. The reference electrode was located at the top of the head, in between the CZ and CPZ electrodes. The impedance of each electrode was lowered to below 5 kΩ before each session began. Data were collected at 1kHz and low-pass filtered under 200 Hz with a notch filter at 60 Hz. The BCI2000 software program was used to process the online data and display the cursor control [35]. Data packets were processed every 40ms to update the cursor position.

First, the data from selected channels were spatial filtered with a small surface Laplacian filter using the four surrounding electrodes for each electrode of interest. Next, an Autoregressive spectral estimation model was used to extract the band power from a 3 Hz bin centered around 12 Hz, here referred to as the alpha power of the signal [36]. The alpha powers of selected channels were used in a linear classifier to produce a control signal for the cursor (2 control signals for 2D tasks, one for each axis). The linear classifier used for each task/axis pair are displayed in Table 1. Finally, the past 30 seconds of classifier outputs are stored in a buffer which is used to estimate the mean and standard deviation of the output signal. These parameters are used to normalize the classifier output to zero mean and unit standard deviation. This final normalized output is used as the cursor velocity.

**TABLE I.** Selected channels and classifiers for Tasks

| Task/Axis | Linear Classifier |
| --- | --- |
| MI-Horizontal | C4-C3 |
| MI-Vertical | -C3-C4 |
| OSA-Horizontal | P3-P4 |
| OSA-Vertical | -PZ-POZ |
| MI+OSA-Horizontal | C4+P4-C3-P3 |
| MI+OSA-Vertical | -C3-C4-PZ-POZ |

For horizontal axes, positive values map to the right direction, while negative values map to the left. For vertical axes, positive values correspond to upwards while negative values result in downwards movement.

### C. BCI Trial Structure

To investigate the effects of combining MI and OSA together, we needed a BCI task that could be controlled with both methods in a similar manner to produce the same output. Virtual cursor control was chosen because it is well established, readily implementable, and intuitive for first time subjects. In addition, the instructions and output are similar between the two tasks (e.g. imagine left hand / look and focus on the left target to move left). Figure 2a displays the trial structure we used for the virtual cursor control task.

**Figure 2.**
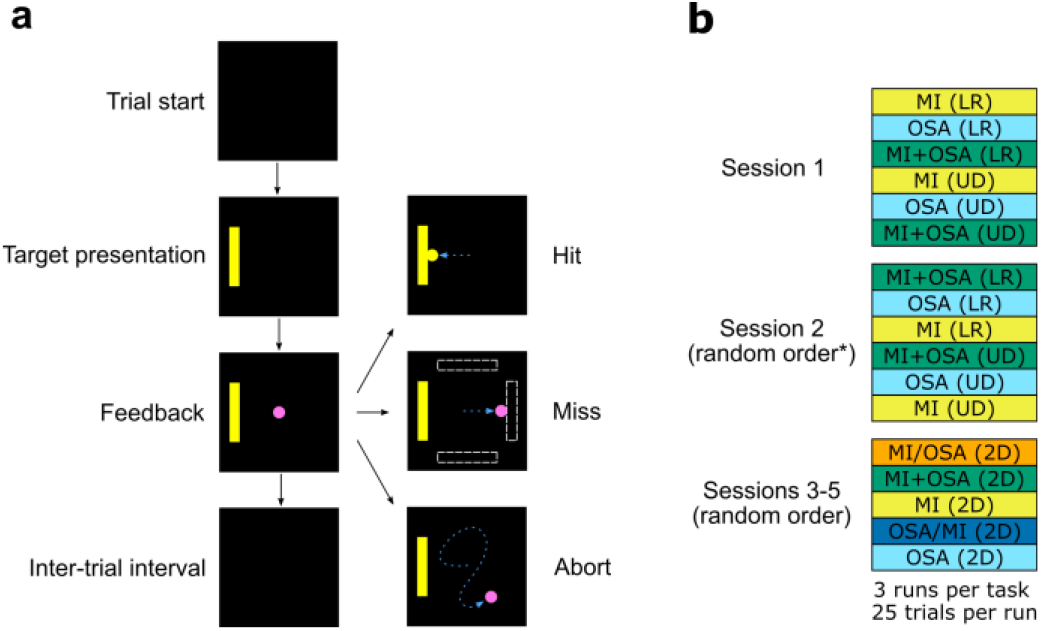
Experiment design of trials and sessions. A) Diagram of BCI trial structure. Each trial begins with a blank screen during the trial start / inter-trial interval. Next, the target, a large yellow rectangle, appeared on either the left, right, top, or bottom of the screen directing subjects where to try to move the cursor. After 1s, a pink ball, the cursor, appeared in the center of the screen which subjects were able to move using the BCI task. This portion of the trial, called the Feedback phase, lasted up to 6 seconds or until a target was hit. If the visible target was hit, the trial ends in a hit, if one of the other invisible targets was hit, the trial ended in a miss, and if nothing was hit in the 6 second time period the trial ended in an abort. After a short inter-trial interval, the next trial began. B) Diagram of the session layout for the experiment. Each subject performed 5 sessions of BCI. During the first two sessions, subjects only performed one-dimensional BCI, either left-right (LR) or up-down (UD). During the final three sessions subjects only performed two-dimensional (2D) BCI tasks. The order of the tasks in the first session was preset for every subject, while the task orders in each of the following 4 sessions were randomized for each subject. *The LR tasks always came before the UD tasks in Session 2, though the order of MI, OSA, and MIOSA was randomized for each subject.

Each BCI trial consists of several phases, beginning with a blank screen. First, a yellow target rectangle appears on either the left side, right side, top, or bottom of the screen. This indicates which direction the subject should attempt to perform to move the cursor. After 2 seconds, the cursor, a small pink circle, appears in the center of the screen and the feedback phase begins. The subject then has up to 6 seconds to move the cursor to hit the visible target, or one of the other 3 targets which remain invisible on the screen. If the subject moves the cursor to hit the visible target, that trial is recorded as a hit. If they hit one of the invisible targets on the other ends of the screen, the trial is a miss. If no target is hit at the end of 6 seconds, the trial is recorded as an abort trial. After the feedback phase, there is a 1 second post-feedback phase where the target and cursor remain frozen on the screen, then a 1 second inter-trial interval phase with a blank screen before a new trial begins. Each BCI run consists of 25 trials of the same task with a randomized order of targets.

### D. Experimental Protocol

Cursor control is commonly performed in either one dimension or two dimensions at once. For the one-dimension tasks, Left-Right (LR) and Up-Down (UD), the cursor is only able to move along the horizontal or vertical axis respectively. These two tasks are intuitive, and some subjects can obtain high performance without much training which makes them good tasks for subjects without any prior BCI experiments. However, some subjects can obtain very good performances with MI or OSA alone (>90% PVC) and since there is a maximum performance at 100% PVC, there is a concern that we would not be able to discern any possible performance improvements form combining MI and OSA (MI+OSA) for subjects who were already performing close to the maximum.

In addition to LR and UD cursor control tasks, two-dimensional control is another well-established BCI task [26], [37]. Here, the cursor can move along both the horizontal and vertical directions simultaneously. This task is inherently more challenging to control than moving in only one direction, which makes it tougher for new BCI users but also potentially a better method for comparing the effects of different control methods by having a higher performance ceiling. Since 2D control is more difficult and subjects can improve over their first few sessions of MI due to a learning effect [20], it can be beneficial for subjects to perform a few sessions of 1D control before moving on to the more challenging 2D task.

We used the study design shown in Figure 2b to be able to analyze 2D cursor control results while also allowing for new subjects to acclimate to the more difficult task by starting with 1D tasks first. Here, subjects performed five total sessions of BCI experiments. During the first two sessions, subjects only performed LR and UD cursor control tasks. This period allowed subjects without any previous BCI experience to train with the easier cursor control tasks before the more challenging 2D setup. The final three sessions consisted of only 2D tasks. Three runs of each BCI task were completed before moving onto the next task, and each run consisted of 25 individual trials. The order of tasks in the first session was the same for every subject, to make sure that subjects with no prior BCI experience could perform both MI and OSA individually before being asked to combine the two together. After the first session, the order of tasks was randomized for every subject individually to eliminate any effects that might occur due to the task order. Although the task order was randomized for the second 1D session, all the LR tasks were always performed before the UD tasks for coherence.

To investigate the effects of combining MI and OSA, we looked at three main tasks: MI alone, OSA alone, and MI and OSA together (MI+OSA). These tasks were performed in both one and two dimensions. In addition, we also wanted to study the effects of using different tasks to achieve different outputs within the same run. To accomplish this, we also ran two new tasks: MI/OSA and OSA/MI. For these tasks, subjects used MI to control the cursor along one axis (either horizontal or vertical) and OSA to control the other. The task before the “/” in the label identifies which task controls the horizontal dimension, while the last task controls the vertical (ex. MI/OSA means MI moves the cursor left and right, while OSA moves the cursor up and down). These two tasks were only performed in 2D sessions since they require two different axes to move upon. Although performing MI and OSA at the same time to move in different directions has been done in previous work, [24] including these tasks in the study design allows us to explicitly compare their performance to other tasks. Figure 3 illustrates the differences between each task, and what the subjects were instructed to do to move in each dimension.

**Figure 3.**
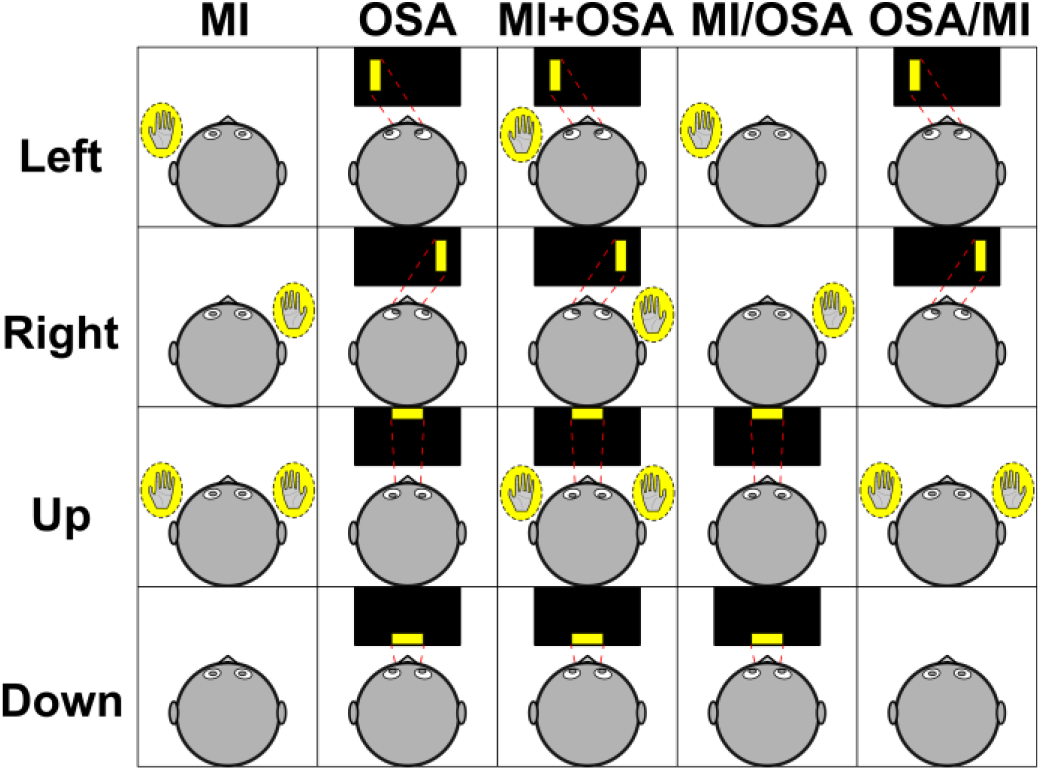
Visualization of MI, OSA, and MI+OSA tasks. Different combinations of hand motor imagery and OSA used to control the BCI cursor in various tasks. Subjects performed motor imagery of either the left hand, right hand, both, or neither. For tasks involving OSA, subjects performed OSA to a target on either the left, right, top, or bottom of the screen. For tasks that did not involve OSA, subjects were not given any restrictions on where to look.

### E. Statistical Analysis

Behavioral performance results were measured using Percent Valid Correct (PVC), a commonly used BCI performance metric. PVC is calculated as the number of hit trials divided by the number of hit and miss trials together (non-abort trials):

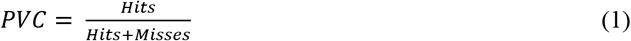

All analysis was performed using the R software environment [38] and the MATLAB toolbox: Fieldtrip. Statistical comparisons were performed using the Wilcoxon Signed Rank test [39], and adjusted for multiple comparisons using the Holm method [40], [41] unless noted otherwise.

## III. RESULTS

### A. Group Level Performance

Figure 4 shows the group level BCI performances for each task as PVC values. Looking at the one-dimension results from the first two sessions (Figure 4 panels a and b), MI+OSA has a higher mean PVC than either MI or OSA in the LR task, but a lower mean than OSA alone for the UD task. After statistical comparison, MI+OSA is only significantly greater than MI in either task. The height of each boxplot shows that subjects have a large variance in performance for most tasks. Some subjects can reach near perfect control with PVC close to 100%, while other subjects do not display any control at all and perform at or below chance level (50% for 1D tasks). The exception to this is the OSA alone control method in the UD task, where subjects achieved a higher performance with a lower variance. In the UD case, MI+OSA performed in between the results of OSA and MI alone.

**Figure 4.**
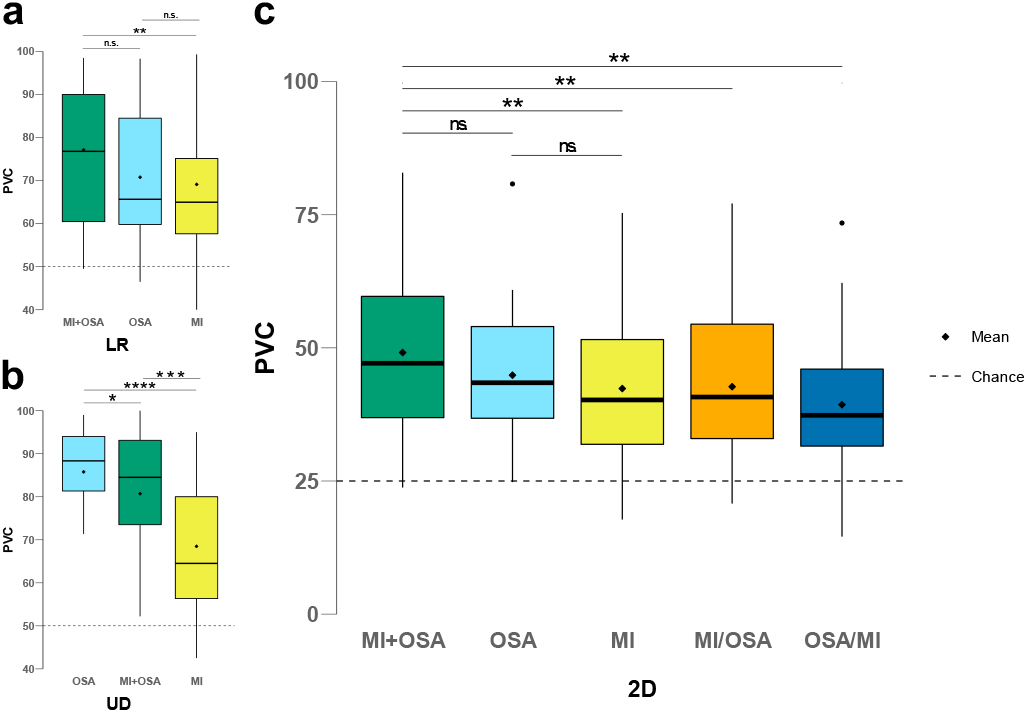
Group level BCI performance. Boxplots showing BCI performance in PVC for each task in one and two dimensions. Statistical testing was limited to comparing MI+OSA to each other task, and MI vs. OSA for the 2D case. A) BCI performance for horizontal (Left-Right) tasks. MI+OSA performs greater than MI alone, but not OSA alone. B) BCI performance for vertical (Up-Down) tasks. MI+OSA again performs greater than MI alone, but not OSA alone. In addition, OSA is shown to substantially outperform MI in the UD task. C) BCI performance for the five 2D tasks. MI+OSA outperforms each method except for OSA alone. OSA is also shown to not outperform MI alone. Statistical testing was performed using the one-tailed Wilcoxon signed rank test and adjusted for multiple comparisons using the Holm method. (n.s.: p>0.05, *: p<0.05, **: p<0.01, ***: p<0.001)

The 2D runs (Figure 4c) show a similar trend as in the LR task. Here, the mean PVC of MI+OSA runs is again higher than the rest of the tasks but does not reach statistical significance against the OSA alone method. The OSA alone method was also not found to be significantly greater than the MI alone method across subjects. The split methods (MI/OSA and OSA/MI) achieved similar performance to the individual methods as well. Subjects achieved worse average performance in the 2D tasks compared to the 1D tasks, which is expected due to the increased complexity, but there are still large variances in performance across subjects. Again, some subjects performed at or below chance level while other achieved relatively high PVC values.

Next, we looked at individual subjects’ behavioral performance data across the different tasks. Figure 5a shows the PVC scores for each task with subjects’ individual performances linked together. From this view, there do not appear to be any clear trends across the tasks. Some subjects perform better using MI alone, while others perform better with OSA alone. Additionally, both of these two groups have some subjects that benefit and some who perform worse when using the combined method. However, this view of the data does suggest that subjects who can only achieve PVCs near or below chance level for one task appear to perform poorly across all tasks.

**Figure 5.**
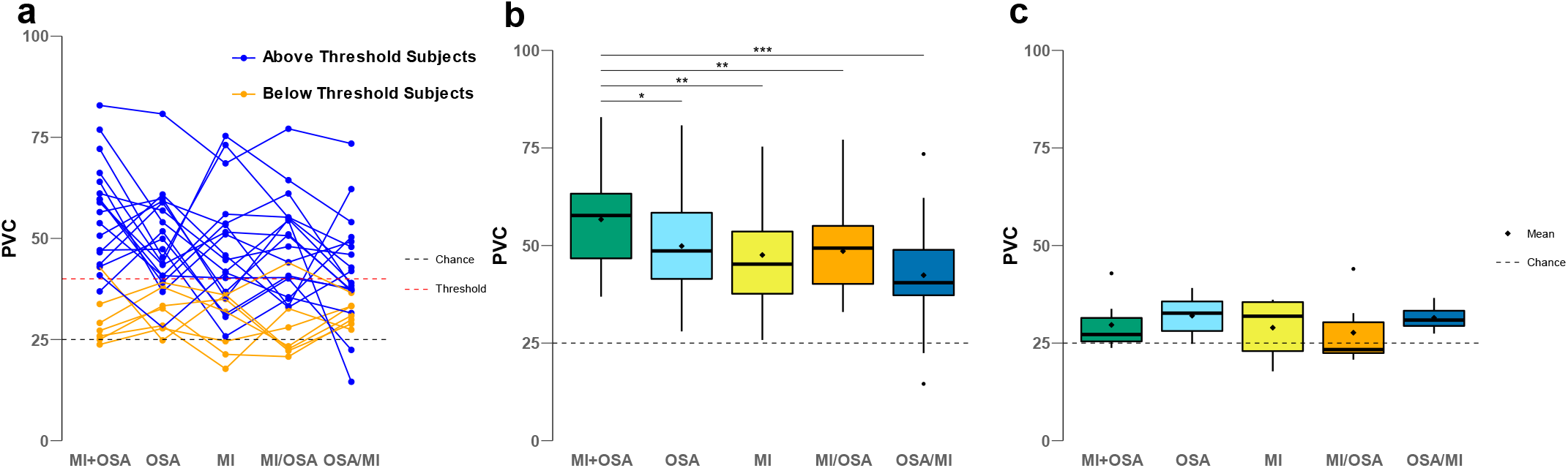
High and low performing sample results. BCI performance for two groups of subjects split by performance in MI only and OSA only. Subjects who obtain an average performance above 40 PVC in either MI alone or OSA alone were placed in the “Above Threshold Subjects” group. The remaining subjects were placed in the “Below Threshold Subjects” group. A) Line plot illustrating individual subjects’ PVC throughout each of the tasks. Subjects that performed lower in either MI or OSA tended to perform poorly in each of the tasks and did not benefit much from combining the methods. B) Boxplot of BCI performance for the subjects above the performance threshold. MI+OSA outperforms each other method for this group of subjects. C) Boxplot of BCI performance for the subjects below the performance threshold. There were no significant differences found among any of the tasks. Statistical testing was performed using the one-tailed Wilcoxon signed rank test for panel B and the two-tailed Wilcoxon signed rank test for panel C. P-values were adjusted for multiple comparisons using the Holm method. (n.s.: p>0.05, *: p<0.05, **: p<0.01, ***: p<0.001)

To study the differences between these lower performing subjects and others who were able to achieve high PVCs, we split the subjects into two groups based on their performance in the MI only and OSA only tasks. Previously, 40 PVC has been considered to be a rough threshold for a subject to be considered BCI proficient.[24] Scores under this threshold indicate that the subject was not able to gain any meaningful control of the BCI system. For this analysis, we used 40 PVC as a threshold to split the subjects into two groups. Subjects who were able to achieve an average PVC above 40 in either MI alone or OSA alone were included in the “Above Threshold” group, while the remaining subjects were included in the “Below Threshold” group. After this split, the difference between MI+OSA and the rest of the tasks was even larger and reached significance against each other task for the group above the threshold. (Figure 5b). We also performed this analysis using two similar performance thresholds (35 and 45 PVC) and found a similar overall trend among the tasks (See Supplementary Figure 1). However, the difference between MI+OSA and OSA was not statistically significant in these cases (p=0.06 and p=0.072 for 35 and 45 PVC respectively). For the lower performing subject group, all the tasks reached similar performance and none of the methods were found to be statistically different than any of the other methods. This was also true for the 35 and 45 PVC thresholds as well. These results provide further evidence that lower performing subjects tend to perform poorly across all the available tasks.

### B. Best Individual Control Method

Comparing the group level performance between different tasks gives a general idea of how well subjects perform using each method. However, since the task of interest (MI+OSA) is a combination of the other two tasks, it may be a more interesting analysis to compare the performance of this method against whichever of the two tasks is better for an individual subject. This comparison illustrates whether combining the two methods allows subjects to achieve better performance than using an individual method, or if combining the two leads to performances somewhere between the individual methods.

For each subject, we determined which method between MI alone or OSA alone allowed that subject to achieve a higher average PVC. This method was then selected as that subject’s “best” method. Comparing MI+OSA and each subject’s best method shows that MI+OSA achieves about the same performance on the group level (Figure 6a). Looking more closely at each individual subject’s average performance, we see that the difference between MI+OSA and each subject’s best method varies greatly across subjects (Figure 6b). This difference ranges from MI+OSA being 23% higher than the best individual method, down to 15.3% lower than the best method.

**Figure 6.**
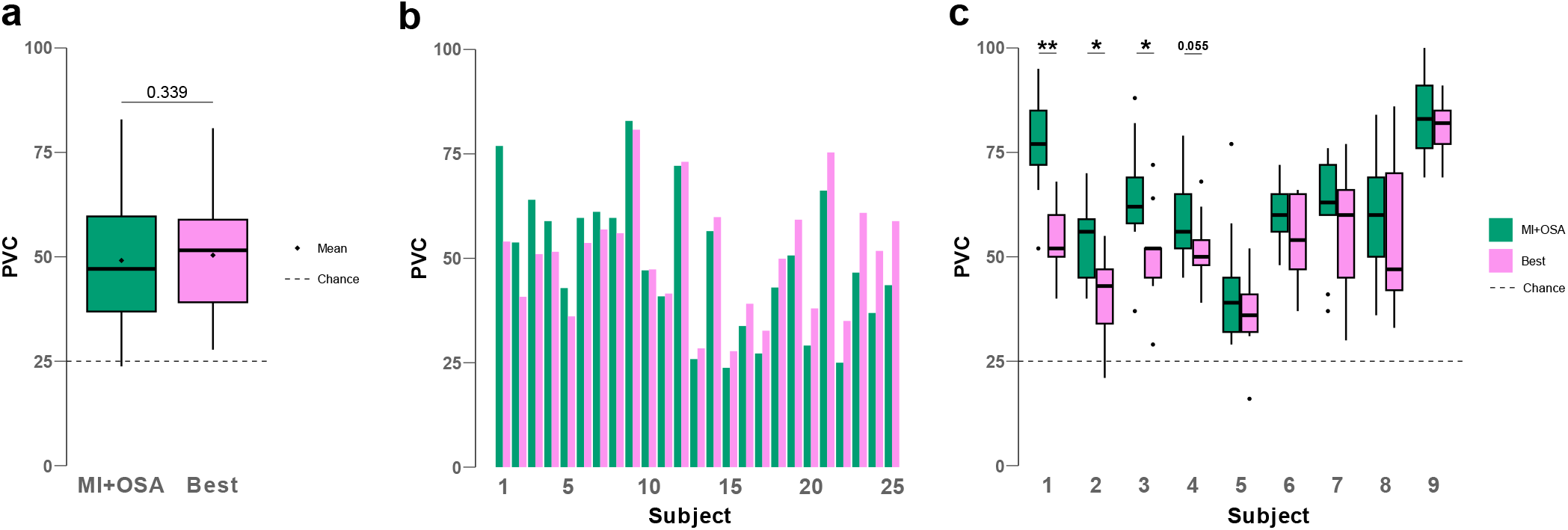
Subject level BCI performance. Two-dimensional BCI performance of MI+OSA compared with each subject’s individual best control method between MI only and OSA only. A) Boxplots comparing the PVC of MI+OSA with each subject’s best method. MI+OSA was found to not be significantly greater than each subject’s best method. B) Bar chart showing each individual subject’s average 2D MI+OSA PVC across 9 runs (green) and average 2D PVC of their best control method (pink). The subjects were sorted by the difference between these two values, with the most positive difference between MI+OSA and Best on the left, and the most negative on the right. C) Individual subject boxplots for MI+OSA and best control. Each boxplot consisted of 9 data points, one for each run of 2D BCI for the task. Only the subjects who performed better in MI+OSA than their best individual control method are shown. Statistical testing was performed using the one-tailed Wilcoxon signed rank test (n.s.: p>0.05, *: p<0.05, **: p<0.01, ***:p<0.001). Note: Adjustment for multiple comparisons was not performed due to the larger number of potential comparisons and the intent of studying each subject individually for this analysis. Applying the Holm adjustment method, only the first subject would remain significant with p=0.0101.

Similar plots as Figure 6b were made to compare MI+OSA to MI alone and OSA alone (See Supplementary Figures 2-4). These plots illustrate that 17 out of 25 subjects had a higher average MI+OSA performance compared to MI alone, and 12 out of 25 performed better with MI+OSA compared with OSA alone.

Figure 6c shows box plots of individual run PVC scores for the 9 subjects who had a higher average MI+OSA PVC compared to their best individual method. Across all 3 sessions, each subject performed 9 runs of MI+OSA and 9 runs of their best method, so each boxplot was constructed from 9 data points. The results show that some subjects were able to achieve substantially better performance using the combined MI+OSA method compared to their performance using their best method out of MI or OSA. While this benefit is limited to just a few subjects, it represents a novel improvement in their BCI performance. It should be noted that many of the subjects did not achieve much better performance with the combined method, and a majority even performed worse with it. However, this may not be too surprising, as the differences in MI only and OSA only PVC scores show that BCI task performance is subject-dependent and what works better for some subjects may not work as well for others.

### C. Electrophysiology

Electrophysiological analysis reveals that combining MI and OSA results in the expected combination of features from both individual methods. Figure 7 shows topological maps of the grand average alpha powers across all subjects for each of the control directions. The Motor Imagery maps show the well-established event-related desynchronization (ERD) of alpha power in the sensory-motor cortex (around electrodes C3 and C4) when subjects are performing different types of hand motor imagery. Interestingly, there is also ERD in the posterior cortex, which is similar to the signals produced by OSA. This suggests that subjects also choose to look and focus on the visible target during MI tasks, even when not explicitly instructed to.

**Figure 7.**
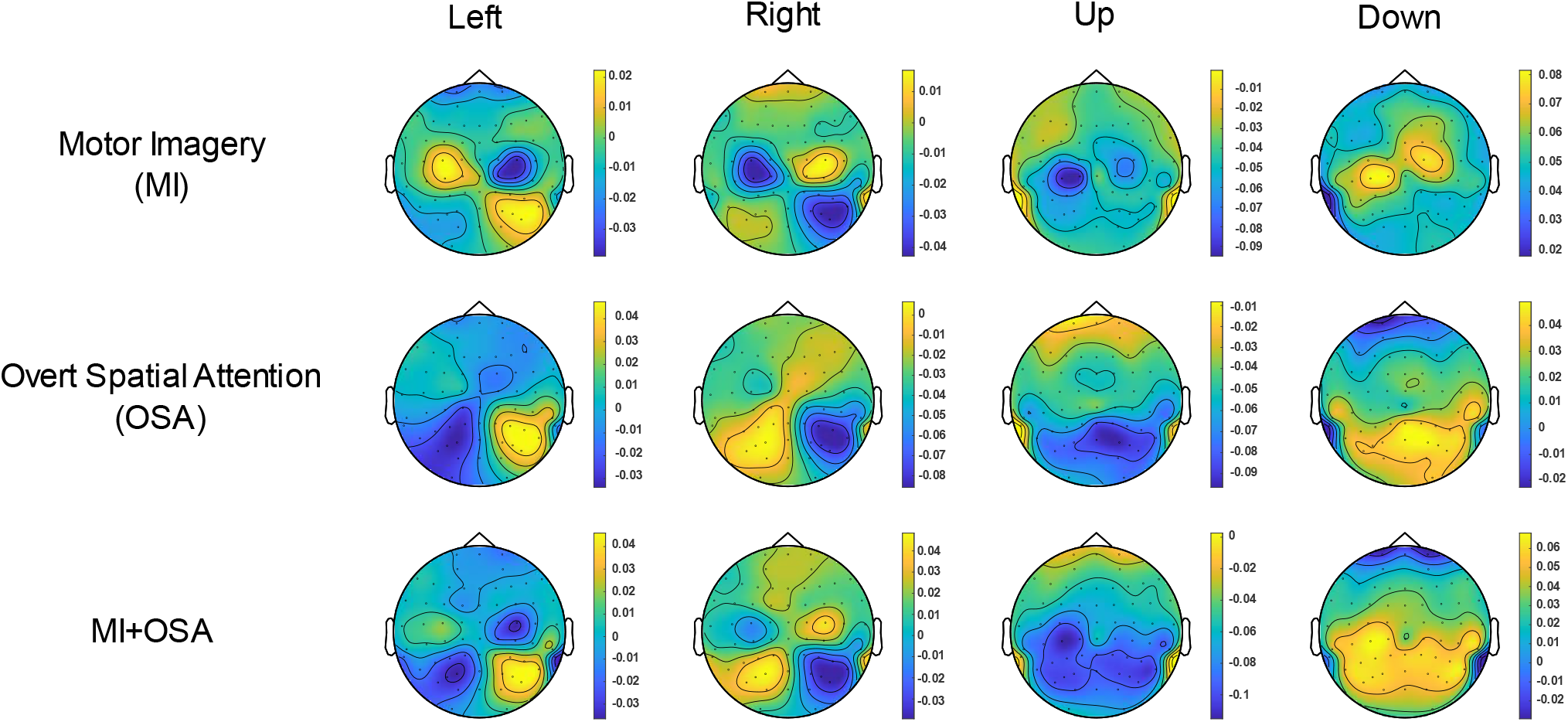
Topographical maps of alpha power changes between tasks. Each plot denotes the relative change in alpha power (8-13 Hz) for each electrode in the montage. Colors between each electrode were interpolated. The topographical maps were obtained from the grand average of alpha band powers across all subjects and runs for the specified control method in 1D tasks. For each target direction (ex. Right), the values are found by comparing the average alpha power of electrodes during those trials against the average alpha power of electrodes in all trials of that 1D task (ex. LR tasks). For example, a lower value in one of the plots in the Up column indicates that performing the Up task leads to lower alpha powers in that cortical area compared to the average alpha power during all UD tasks.

The middle plot shows the relative band powers in the alpha band for OSA tasks. Again, the results follow previous works

[24] and show event related desynchronization in the posterior cortex on the same side as the target for LR tasks. For Up tasks there is ERD in the central posterior cortex. The bottom two maps show relative band powers for the combined MI+OSA tasks. These maps shown a combination of features from both the MI and OSA maps, which is expected from the task definition. In each of the maps from different control directions, there are features from both the Sensory-motor cortex (center band of electrodes, especially C3 and C4) as well as the Posterior cortex (lower electrodes, especially P3, P4, Pz, and POz). This same data is shown in Supplementary Figure 5, using color bars centered around zero. This alternate representation has less contrast between the features but uses blue and yellow strictly for positive and negative differences, which may be helpful for comparing the relative strengths of different features across the maps. These results show that, on a grand average scale, the neural signals that are produced by both MI and OSA individually can also be produced simultaneously in a similar manner.

## IV. DISCUSSION

In this work, we have investigated a new BCI control paradigm, MI+OSA, which integrates two established BCI control methods together simultaneously. The results show that MI+OSA is an effective BCI control method that reached the highest average performance among paradigms studied in the 2D task, and statistically outperformed using MI alone. In addition, OSA alone was found to not be significantly greater than MI in the 2D tasks, while MI+OSA was. This suggests that, on average across all subjects, MI+OSA seems to be the best control method among the studied paradigms, though more evidence is needed to show that the improvement over OSA alone is not due to chance. Our results also show that MI+OSA achieves similar performance to each subject’s best personal paradigm between MI alone and OSA alone, which suggests that combining MI+OSA can lead the decoder to using the best available features between the two individual paradigms. Looking at individual subjects’ performances, we also found that some subjects performed better with MI+OSA compared to either of their MI alone or OSA alone scores. Since MI+OSA is a combination of these two paradigms, this shows that, for some subjects, the decoder is able to take advantage of features from both paradigms to achieve a higher performance than using either individual method alone.

The group level results in Figure 4 show that the MI+OSA task reached the highest average PVC across all subjects for the LR and 2D tasks. However, after statistical testing, the difference between MI+OSA and OSA in both cases was not found to be statistically significant (p<0.05). Although there is a numeric difference between the mean performance in both tasks, this difference is relatively small compared to the large variance between subjects’ BCI performances in both LR and 2D tasks. While twenty-five subjects is not a small sample size for a BCI study, many more would likely need to be studied to achieve the statistical power necessary to conclude that the numerical difference in means between MI+OSA and OSA is not due to chance.

In the UD case, OSA had the highest average PVC over MI+OSA and MI alone. Since MI+OSA is a combination of the other two methods, its performance landing in between the two individual techniques suggests that instead of combining features from both tasks, the features from the lower performing task (MI in this case) may be reducing the decoder’s ability to make correct decisions using the features from the higher performing task (OSA in this case). The difference in this result compared to the LR and 2D cases, may be due to the large performance difference between MI alone and OSA alone in the UD case, which is not present in the LR and 2D tasks.

The MI+OSA topographical maps in Figure 7 shows that there is a widespread ERD in the posterior areas of the cortex in Up MI+OSA trials compared to Down trials. Some of these areas do not seem to be activated in either MI alone or OSA alone UD tasks, so there may be additional features that could be decoded in MI+OSA UD trials that could improve performance. Overall, combining MI+OSA does not seem to substantially benefit subjects compared to using OSA alone on the group level.

Group level statistics provide an intuitive overview of how each method compares across all subjects together. However, each subject has their own features that arise from using the different methods, making BCI decoding inherently a subject-specific challenge. The traces in Figure 5a. demonstrate this point by showing that subjects follow various patterns across the different tasks. Here, it is not clear that any one method is better than another because each subject’s performance varies across the different tasks. For example, some subjects performed better with MI over OSA, while others had their best performance with OSA, so it is hard to conclude that either one is a better method overall.

Figure 5a also shows that subjects who perform poorly or near chance level (25% PVC) tend to perform poorly across all of the tasks. By splitting subjects by their PVC in the MI only and OSA only methods, we found that there was a larger positive effect of using MI+OSA compared to the other tasks for the subjects above the performance threshold (Figure 5b). Although at first this suggests that MI+OSA is more beneficial for subjects with better individual MI and OSA performances, some of the best performing subjects still did not benefit from MI+OSA and even performed worse with it. Figure 5c illustrates that the lower performing subjects performed nearly as bad in each task across the board. Therefore, the results suggest that subjects do not benefit from MI+OSA when they have low performance with the individual methods, rather than suggesting that higher performing subjects benefit from it more.

Since MI+OSA is a combination of two individual methods, an argument may be made that it is only worth using if it performs better than either method alone. Otherwise, subjects could just use their best individual method to obtain better performance, since they must be able to use both MI and OSA for the BCI application in order to use MI+OSA in the first place. Therefore, instead of comparing MI+OSA to MI and OSA it may be a more informative analysis to compare MI+OSA to each subject’s best individual method. The results in Figure 6 illustrate this comparison through multiple analyses. Figure 6a shows that subjects achieve about the same performance using MI+OSA compared to their best individual method on the group level. This suggests that, at least on average, combining MI and OSA still leads to the simple online decoder using the best available features from between the two methods.

Figure 6b shows the various performance differences between MI+OSA and the best individual method for each subject. These results again emphasize the subject-specific nature of BCI performance. Some subjects benefitted substantially from MI+OSA, some performed much worse with it, while most subjects achieved similar performance with their best individual method. The results in Figure 6c take a closer look at each individual subject’s performances and show that about 4 out of the 25 subjects were able to achieve considerably better performance using the combined method. With these results, MI+OSA may not appear to be a useful method when looking at the entire study population together since only a small portion of the subjects benefitted from it. However, the goal for each BCI user is to obtain the best control of the system that is possible for them. From this point of view, MI+OSA is valuable as an additional control method option that could significantly improve BCI control, even if only for some subjects. More sophisticated decoders could have the potential to increase the number of subjects that could benefit from MI+OSA, and even increase the performance gain.

Figure 7 shows the grand average topographical maps for 1D tasks using each method. These plots serve two functions. First, they confirm that subjects performed the correct tasks for MI and OSA by producing similar electrophysiological features as in previous studies. As expected, MI produced ERD in the sensorimotor areas and OSA showed ERD around the posterior cortex. The MI plots also show some activity in the posterior cortex, similar to what is expected for the OSA task. One explanation for this is that subjects were free to look and focus wherever they like during the MI tasks, and it may be natural for subjects to look and focus on the target while performing the corresponding MI. Therefore, they may have unintentionally been performing MI+OSA during MI only tasks, even when not specifically asked to. The major difference between the MI+OSA and MI tasks in this case is that the online decoder was only focused on sensorimotor electrodes for MI (C3 and C4), while it used both sensorimotor and posterior cortex electrodes for MI+OSA (C3, C4, P3, P4, Pz, and POz). The second function is to investigate the cortical activity produced by the MI+OSA task. The MI+OSA topographical maps show a combination of the features for MI and OSA. Since this task is a combination of the two individual tasks, it is not surprising that the features are also a combination of features from the individual tasks. One important result from these maps is that both MI and OSA features can be activated simultaneously. This suggests that other independent BCI tasks may also be combined simultaneously and decoded together as well. Otherwise, these plots reinforce the expected electrophysiological features that are expected from MI, OSA, and the combination of the two.

A limitation of this study is the simplicity of the traditional online decoder. Although the topographical maps in Figure 7 show that MI+OSA results in potentially decodable features in more electrodes across the cortex, the decoder used in this analysis may not have been able to take advantage of these additional features. The simple linear classifier used in the online experiments weighed the ERD in each of the electrodes of interest evenly. Therefore, if a subject produced strong ERD in an electrode using either MI or OSA, the features in this electrode would still overshadow the others in MI+OSA. While this may have helped some subjects’ online performance by continuing to use the strongest features available, more sophisticated decoders may be able to take advantage of smaller ERD/ERS features that occur in other electrodes to improve performance even more. In addition, the simple linear classifier used for the online experiments used electrodes and frequency bands that were preselected before the study based upon prior knowledge of MI and OSA features and were the same for each subject studied. This simple decoding strategy was straightforward to implement and allowed us to readily compare different subjects’ performances. However, looking at individual subjects’ topographical maps for the various tasks, there is noticeable variation in subjects’ ERD patterns. This means that subjects would almost certainly benefit from a decoding strategy that takes advantage of subject-specific features. Incorporating these subject-specific decoders may allow some subjects who performed at or near chance level to achieve more effective BCI control and could potentially allow more subjects to benefit from MI+OSA.

Future work could be done to further characterize any potential benefits of using MI+OSA compared to each individual method. Using more sophisticated decoders for further offline analysis may reveal additional features from MI+OSA or provide a more consistent control signal. Although this was not the case in the online results for most subjects, more advanced signal processing may allow subjects to achieve higher performances with MI+OSA, such as suggested by recent studies using electrophysiological source imaging [3], [15], [42] and deep learning [19], [21], [22].

## V. CONCLUSIONS

The results from this study show that combining two established BCI control methods, MI and OSA, results in a hybrid method that achieves similar results to each subject’s best individual method, but also allows some subjects to obtain their best BCI performance. This work shows the potential of combining BCI control methods to improve performance and reliability of BCI devices, as well as the need for developing more BCI control options for users. Since different users perform best using different control methods, providing more BCI control options may allow more potential users to reach effective BCI control. Overall, improving the performance and reliability of these non-invasive BCI devices will expand their potential clinical uses, and may help to restore or replace motor functions for patients in need and can even provide direct brain-to-device communication in the general population.

