## Supplementary Materials for "Integrating simultaneous motor imagery and spatial attention for EEG-BCI control"

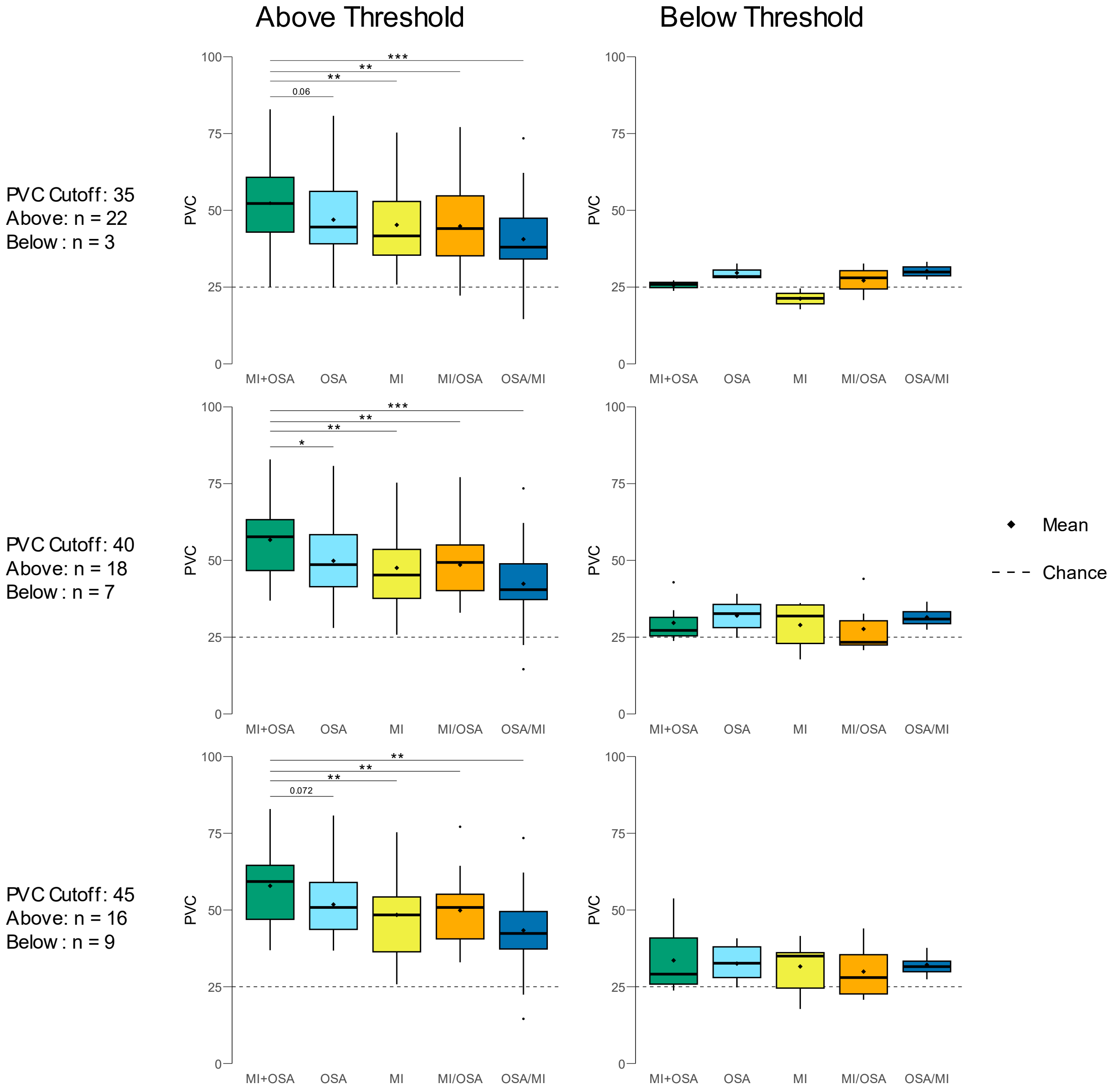

**Supplementary Figure 1. Different splits based on subject performance.** To investigate the differences between high and low performing subjects, we split the total study population based on their BCI performance in MI alone and OSA alone compared to a performance threshold. (See Figure 5 in the main article). Here, the same analysis and plots were generated using slightly different thresholds. From top to bottom, the thresholds were 35, 40, and 45 PVC. The plots on the left side include subjects who reached an average PVC greater than the threshold using either MI alone or OSA alone, while the plots on the right include subjects who did not reach this threshold.

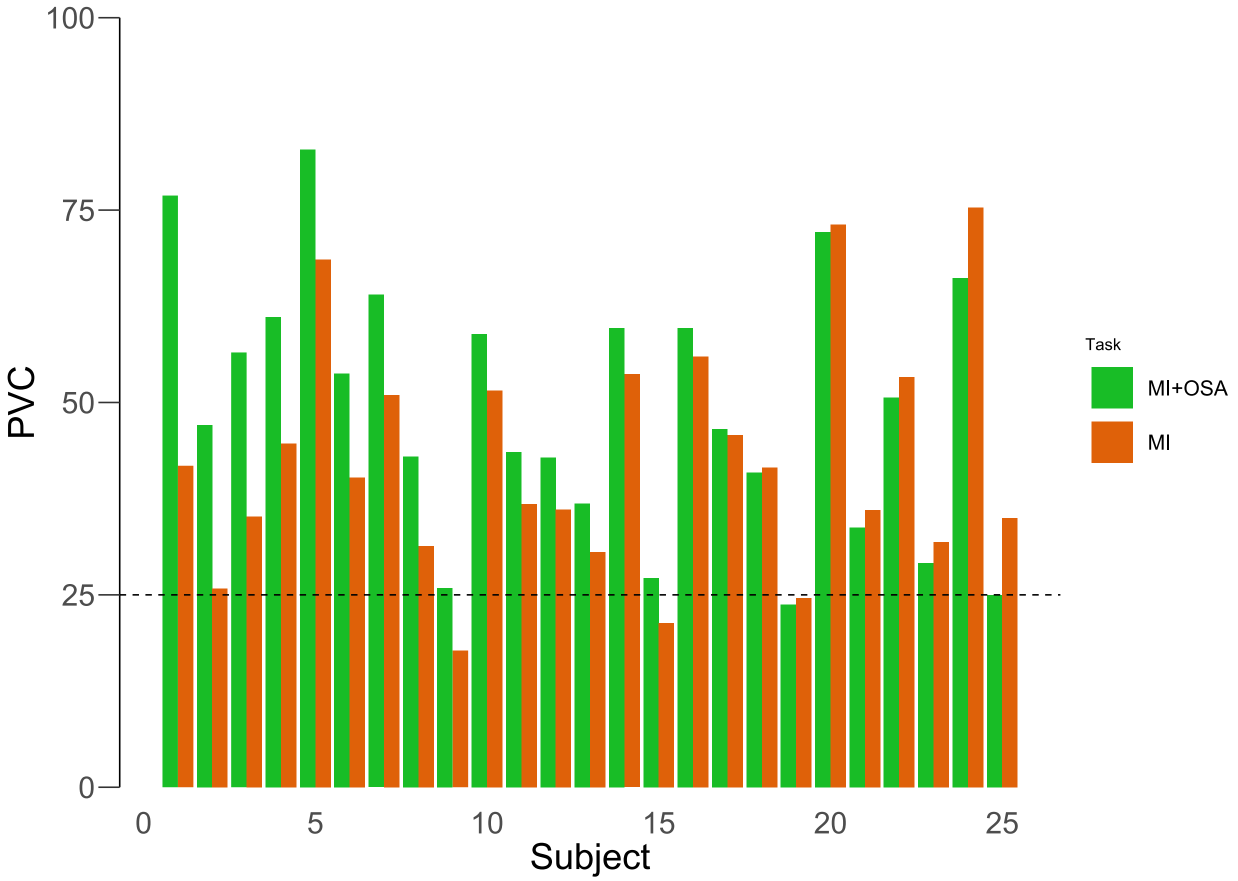


**Supplementary Figure 2. Individual subject MI+OSA vs MI performances.** Each pair of bars shows the average 2D MI+OSA and MI scores for individual subjects. The subjects are sorted in order of the difference between their MI+OSA and MI performances, with the subjects on the left having the largest MI+OSA compared to MI, and the subjects on right having the lowest MI+OSA compared to their MI performance. From the plot, we can see that 17 out of 25 subjects performed better with MI+OSA compared to MI alone.

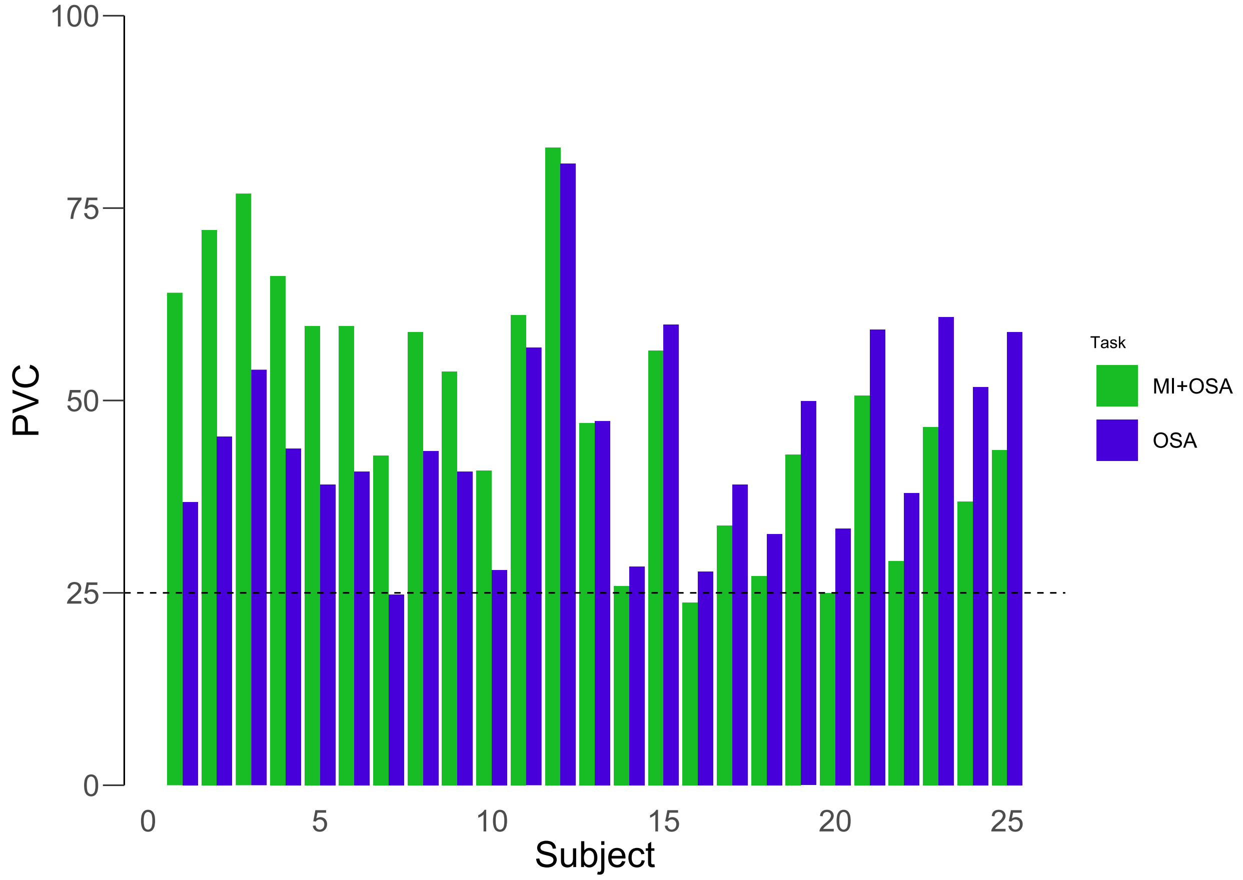


**Supplementary Figure 3. Individual subject MI+OSA vs OSA performances.** Same analysis as Supplementary Figure 2 with OSA instead of MI. Subjects were again sorted by the difference between their MI+OSA and OSA performances, with the larger MI+OSA performance on the left. The plot shows that 12 out of 25 subjects performed better with MI+OSA compared to OSA alone.

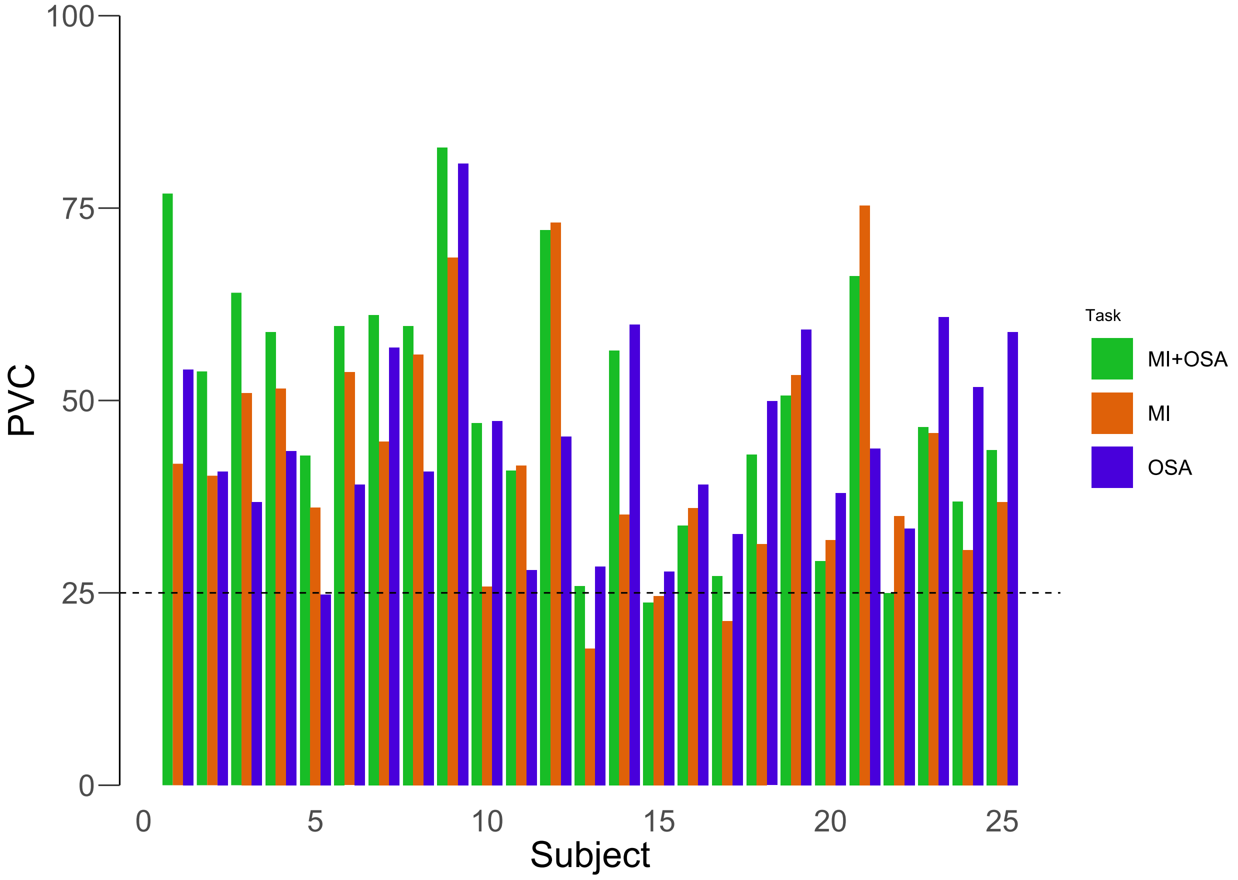


**Supplementary Figure 4. Individual subject MI+OSA, MI, and OSA performances.** The same data from Supplementary Figures 2 and 3 compiled into one plot. Subjects were ordered in the same manner as Figure 6 in the main text.

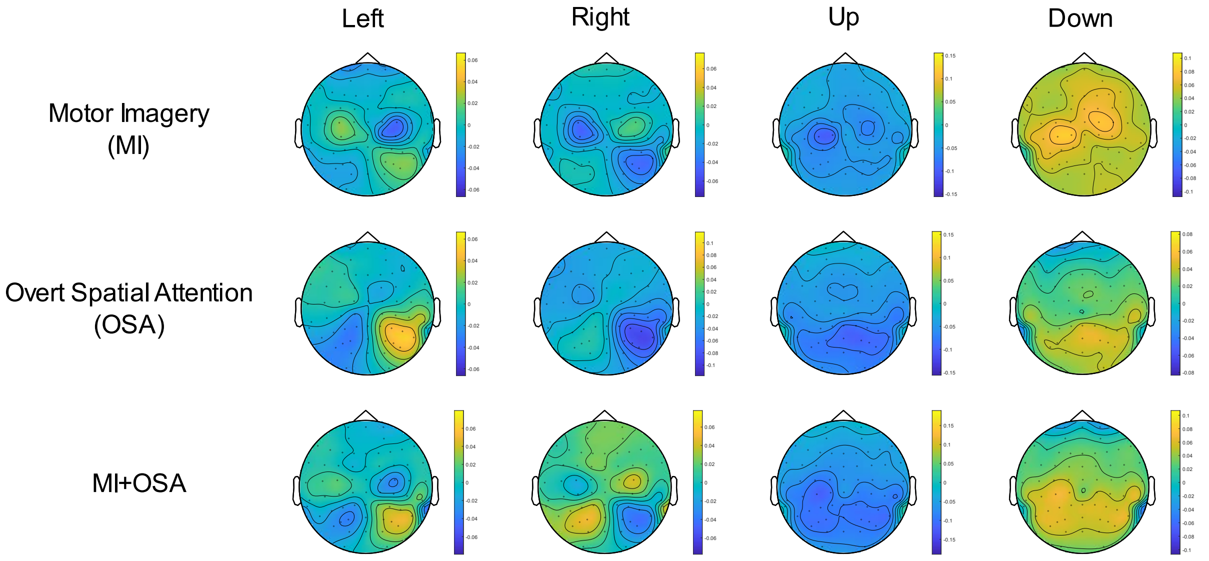


**Supplementary Figure 5. Topographical plots with centered color bars.** The topographical plots here are made from the same data as in Figure 7 in the main text, but the color bars have been shifted and scaled to be centered around zero and extend only up to the largest magnitude in the plot. This alternative visualization may be useful as the blue and yellow colors strictly represent positive and negative differences between the control direction tasks (ex. Right tasks) and the average alpha powers among all tasks in the 1D experiments (ex. LR). For example, both the MI and MI+OSA plots for the left direction have positive yellow features around P4. However, the representation here makes it clear that this feature is stronger in MI+OSA compared to MI alone. While this is also true in the Figure 7 of the main text, it is less clear due to each color bar being scaled individually for each topographical map.
